# Limited horizontal transmission of an obligate, free-living bacterial symbiont

**DOI:** 10.64898/2026.06.25.734684

**Authors:** Liam T. Sullivan, Suzanne E. Kelly, Martha S. Hunter

## Abstract

Nutritional symbionts can be essential for their animal hosts. The bacterial symbiont of the leaffooted bug, *Leptoglossus zonatus*, *Caballeronia*, is acquired from the environment each generation in the 2^nd^ instar. The symbiont is critical for *L. zonatus*: aposymbiotic bugs are unable to reproduce. We hypothesized that symbiotic bugs excrete *Caballeronia* where juveniles might find and consume them. We inoculated *L. zonatus* with GFP-labelled *Caballeronia* and examined feces of each life stage. We found that *Caballeronia* is excreted almost exclusively in the adult stage. We then asked if 2^nd^ instar nymphs could acquire *Caballeronia* from feces. Nymphs were provided with a) feces from adults fed GFP-labelled *Caballeronia*, b) GFP-*Caballeronia* in culture, or c) water only. We found that feces-fed bugs had similar rates of symbiont acquisition to those fed *Caballeronia* in culture, indicating that feces can be a source of *Caballeronia* for *L. zonatus*. However, compared to culture-fed individuals, bugs fed feces had reduced survivorship and required longer to develop, and surviving adults had reduced mass. Bacterial motility assays showed that in contrast to cultured *Caballeronia* cells, *Caballeronia* in feces were non-motile. These results suggest that feces can be a source of *Caballeronia,* at least in some environments. However, transmission mode can influence the success of the offspring.

## Introduction

Microbial symbionts can be critical to the fitness of their animal hosts [1]. Symbionts provide a variety of benefits for their hosts, including the synthesis of limiting nutrients such as essential amino acids and B vitamins (e.g. [2]), recycling nitrogenous waste [3], and digesting complex molecules like lignin [4]. In many cases, these associates are integral to the survival of their host and their maintenance and transmission critical for host fitness. For benefits to be conferred, a host must have a stable means of transmitting a symbiont across generations or among individuals within a community. In insects, obligate symbionts are commonly vertically transmitted from mother to offspring or, as in social insects, horizontally from parents or other conspecific individuals through trophallaxis (anal or oral feeding) or coprophagy (consumption of feces) [5,6]. More unusually, some insects acquire their obligate partners from environmental sources, as in the marine *Vibrio*-squid [7,8] and *Candidatus Endoriftia*-tubeworm symbioses [9]. The transmission success of symbionts that are passed vertically from mother to offspring is often near perfect, as horizontal transmission can be in social insects [10,11]. By contrast, insects that acquire their symbiont from the environment must first locate it. This strategy may present risks, especially in a heterogeneous terrestrial environment [8,9,12].

The arboreal leaffooted bug, *Leptoglossus zonatus* Dallas (Hemiptera: Coreidae) belongs to one of nine families within the Lygaeoidae and Coreiodea superfamilies (Hemiptera: Pentatomomorpha) that associate with free-living soil bacteria of the genus *Caballeronia* (Betaprotobacteria: Burkholderiaceae, formerly *Burkholderia*). First described by Kikuchi and colleagues [13], *Caballeronia* is acquired from the insect’s environment beginning in the 2^nd^ instar and is housed within the specialized midgut 4 region (M4). Here, the growing *Caballeronia* population synthesize essential amino acids and B vitamins [14] and are themselves digested by the insect host. While the reliance on *Caballeronia* is variable among its associated bug host species, the symbiont is obligate for *L. zonatus*. Without it, *L. zonatus* has reduced survivorship and adult weight, and does not reproduce [15,16].

Studies of *Caballeronia* in the model host *Riptortus pedestris* Fabriscius (Hemiptera: Alydidae) have shown a complex and highly specialized interaction. *Caballeronia* is ingested, along with many other microbes, by the 2^nd^ instar nymph [17–23]. To isolate their symbiotic partner, the host bug employs a strict filter, the constricted region (CR), between the midgut 3 (M3) and M4 sections. The CR functions as a physical barrier to non-symbiotic bacteria [24] and can only be passed by *Caballeronia* and allied genera. While *Caballeronia* comprise a comparatively small portion of the soil community, often only ∼10^5^ cells per gram of soil, very few cells are required for successful colonization, only ∼80 cells [25]. The colonizing bacteria wrap their flagella around their cell body and move through the CR lumen in a corkscrew-like movement [26,27]. Only through this mechanism can *Caballeronia* enter the M4 and proliferate. Upon entering the M4, the flagella are lost and the cells take on an ovoid shape, although the host bacteria again may become flagellate when extracted from and grown outside the bug [14]. Curiously, in *R. pedestris,*there is no evidence that *Caballeronia* is released or escapes, at least while the host is alive, and hence there is no transmission from symbiotic parents to aposymbiotic offspring in this species [13]. However, recent studies have demonstrated that the squash bug *Anasa tristus* (Hemiptera: Coreidae) excretes *Caballeronia* in feces, and nymphs orient towards and acquire the symbiont horizontally by ingesting conspecific feces [28]. Similarly, another coreid, *L. phyllopus*, may enrich soil populations of *Caballeronia* through excretion [29]. While the presence of flagella is essential to the colonization of the M4, morphology and flagellate status in the feces of coreid bugs is unknown.

A native to the western United States and South America [30], *L. zonatus* feeds on trees like desert willow [31] as well as orchard crops such as pomegranates, almonds, and citrus [32,33]. Previously we have shown that for *L. zonatus*, *Caballeronia* can be difficult to acquire in the foliage or stems of the tree canopy environment, suggesting that nymphs must undertake potentially treacherous journeys from canopy to soil to locate their symbiotic partner [34]. Additionally, we found that the timing of symbiont acquisition is restricted to approximately one week following molting to the second instar. After this time, fitness suffers: individuals that experienced a delay of greater than one week before symbiont acquisition had reduced gut morphogenesis as well as delayed development, reduced adult weight, and reduced survivorship [35]. Taken together, should *Caballeronia* be rare or limited in the environment, the risk of acquisition failure and acquisition delay increase. In this context, we explored a potential alternate route for transmission of *Caballeronia* between generations, horizontally from adult feces, which may supplement available environmental *Caballeronia* for juvenile bugs.

In this study we asked 1) can *Caballeronia* be excreted in the feces of different stages of *L. zonatus,* and, if so, 2) can feces serve as a symbiont source for aposymbiotic nymphs? We also asked 3) are *Caballeronia* cells in the feces motile? Bacterial motility suggests a flagellate state, required for transit of the CR and colonization of the M4. In the context of the apparent rarity of successful symbiont acquisition in the canopy environment and the severe cost of acquisition delay or failure for these insects, we asked if horizontal transmission via feces for *L. zonatus* might help mitigate these risks, and how alternate routes of symbiont transmission might impact host performance.

## Methods

### Insect rearing

*Leptoglossus zonatus* were reared in the laboratory at the University of Arizona at 27°C, 16:8 L/D on potted cowpea (*Vigna unguiculata*) plants and fed raw peanuts as described in [34]. Eggs were collected at 24h intervals in Petri dishes to synchronize emergence and development. After hatching, first instar nymphs were given deionized water with 0.05% ascorbic acid (DWA; [13] in 2 ml vials with cotton plugs. Upon reaching the 2^nd^ instar and prior to the start of each experiment, water was withheld for 24h to encourage symbiont ingestion.

### Bacterial culturing and insect inoculation

Green fluorescent protein (GFP) – transformed *Caballeronia* strain LZ049G is a strain in the δ (= Coreoidea) subclade, which currently contains no named species [19,36]. The wild form was isolated from a single *L. zonatus* adult collected in Tucson, Arizona in 2018 at the UA West Campus Agricultural Center (16). The bacterium was transformed using a Tn7 insertion incorporating gentamicin resistance. Bacterial culturing and insect feeding followed methods developed by Kikuchi and colleagues [13,37] with slight modifications. For feedings, bacteria were grown on [38] yeast-glucose (YG) media [37] at room temperature (∼18°C–20°C) and liquid YG media was inoculated from single colonies. Liquid culture was grown at room temperature on a rotary shaker (280 rpm) overnight. Prior to presentation to nymphs, cells were either washed in sterile DI water and resuspended in DI water to ∼ 10,000 CFU/ul (for the adult feces transmission and bacterial motility assays) or standardized to 0.1 OD in fresh YG broth (for the juvenile excretion experiment).

For *L. zonatus* used to determine the presence of *Caballeronia* in adult feces, nymphs in the *Caballeronia*-fed positive control treatment were fed ∼2.5 ml of *Caballeronia* (GFP strain LZ049G) in suspension on nonsterile cotton gauze. After 24 hours, symbiont cultures were removed and replaced with DWA and peanuts.

### Microscopy

Fluorescence microscopy to detect the GFP-labeled symbiont was performed using an Olympus BX53 microscope with a white light source, X-cite 120LED Boost illumination system, 488 wavelength filtercube, and 2× 10x, 40x 60x, and 100x objectives. Images were captured using an Olympus DP74 camera and processed with Olympus cellSens Standard 1.6 software. Image analysis was performed with Fiji, a distribution of ImageJ2 [39].

### Stage-specific symbiont excretion

A cohort of isolated 2^nd^ instar nymphs was generated (N = 70) in 7 replicates of 10 individuals and were kept in deep Petri dishes (25mm H × 100mm W) with peanuts and with deionized water with 0.05% ascorbic acid (DWA) *ad libitum*. All nymphs received fresh LZ049G *Caballeronia* in the early 2^nd^ instar for two consecutive days. At the beginning of each nymphal stadium of the symbiotic nymphs, feces were collected to assess the presence of the symbiont, as described in more detail below. To compensate for slight variation in development time, even-staged cohorts were created by transferring newly molted bugs into new Petri dishes with bugs from other dishes that molted on the same day. When nymphs reached the 5^th^ instar, the number of bugs per dish was reduced to a maximum of 5 to provide adequate space for eclosion. Bugs were reared until they reached adulthood, after which 12 bugs were randomly selected to assess the M4 symbiotic organ for the presence of GFP tagged *Caballeronia* using fluorescence microscopy.

To assess whether feces contained *Caballeronia*, fresh feces were collected once daily during the first 48 hours of each life stage. At this time, insects were transferred into sterile Petri dishes to allow feces to accumulate. For the 2^nd^ instar, feces collection began one day after symbiont feeding concluded. Following each 24hr period, up to five fecal droplets were collected from each replicate. Individual fecal droplets were gently flushed from the Petri dish surface with 10µl of sterile 1x PBS using a micropipette and transferred to YG media containing 50µg/mL each of cycloheximide and gentamicin and allowed to air dry. This inclusion was designed to prevent growth of environmental contaminant bacteria and fungi that were likely to be present in or on the collected droplets. Plates were incubated at room temperature and checked daily until colonies formed or the experiment had ended, at about 4 weeks.

### Symbiont transmission through adult feces

After confirming that adult feces contained *Caballeronia,* we asked if *Caballeronia* could be acquired from adult feces by nymphs and if the age of the excreting bug affected transmission success. To test this, isolated aposymbiotic 2^nd^ instar nymphs were presented with fresh feces collected from a cohort of *L. zonatus* previously inoculated with GFP-labelled *Caballeronia* (LZ049G) and reared to adulthood. A factorial design was used to include the impact of age of the excreting adult on fecal transmission. Nymphs were reared in two temporal blocks, each block with each of 3 treatments presented to 2^nd^ instar nymphs: feces-fed (see collection methods below), *Caballeronia* from culture-fed (for a positive control), or symbiont-free, receiving only water (for a negative control). Temporal blocks were used to determine if the age of the excreting adult bug affected the successful transmission to nymphs. Excreting adults were either young (24-48 hours after eclosion) or mature (2 weeks following eclosion). Each treatment/block was replicated 8 times, with six 2^nd^ instar nymphs per replicate. To collect feces, a cohort of GFP-fed bugs was generated and reared to adulthood. Upon eclosion, bugs were randomly assigned to groups of three and given DWA, cowpea sprigs (*Vigna unguiculata*), and raw organic peanuts to feed upon. For feces collection, adults were placed in clean, clear 473mL Solo brand cups fitted over sterile 100mm Petri dish bases where fresh feces were collected. After feces accumulated for 48 hours, the feces-containing Petri dishes were removed. Aposymbiotic nymphs were introduced into the frass-ladened Petri dishes for the frass-fed treatment. All dishes were gently misted with DI water once per day to keep the feces moist until nymphs reached the third instar, after which bugs were followed throughout development to adulthood.

### Bacterial motility in adult feces

To assess bacterial motility in feces, we collected fresh (<24hr old) liquid feces from *L. zonatus* adults colonized by LZ049G. One µL of frass was transferred to a glass slide and assessed with the fluorescence microscope following methods modified from [40]. Short videos were recorded at ∼200ms exposure (5 frames per second) under 40x magnification. Video footage was then examined, and bacteria were considered motile if cells were observed moving independent of the fluid flow with an average velocity of >4 µm/sec. For examples of motile and nonmotile bacteria, see SI videos 1 and 2, respectively. Five cells were selected from each sample to estimate velocity. Measurements were performed using the Manual Tracking plugin in FIJI [39]. A sample was considered motile if the average velocity of the measured bacteria was >4 µm/sec. To assess the flagellate state of LZ049G *in vitro,* we examined bacteria grown in liquid culture as described above. For *in vivo* conditions, insects were inoculated with *Caballeronia* LZ049G in the 2^nd^ instar, and after eclosion, adult insects were sacrificed and dissected. The M4 was removed and a portion placed in 10µL of sterile 1x PBS, where it was broken open and cells in the surrounding fluid were recorded to assess motility.

### Statistical Analyses

All statistical analyses were performed using R version 4.5.2 [41] and Rstudio [42]. Figures were created using ggplot2 [43].

Adult weight and development time were both initially assessed using a linear mixed model implemented with *glmer* (*lme4* [44]) with age group and treatment as explanatory variables, and replicate included as a random variable. For adult weight, sex was also included as part of the model, as *L. zonatus* females are often larger than males [16]. The inclusion of replicate resulted in singularity of fit for the analysis of adult weight, and the model was simplified to a linear model implemented with base R. Backwards model selection was implemented using *drop1* [41]. Rates of acquisition success and survivorship to adulthood were assessed using a generalized linear mixed binomial model with age group and treatment as explanatory variables and replicate as a random variable. The inclusion of replicate resulted in singularity of fit for the analysis of survivorship, and the model was then simplified to a generalized linear binomial model. Where appropriate, posthoc comparisons were performed using a Tukey’s test implemented with *glht* (*multcomp* [45]).

Bacterial motility and the presence of *Caballeronia* in juvenile frass were analyzed using Fisher’s exact test (*rstatix*, [46]). Motility was assessed as the proportion of samples (videos) with motile bacteria across treatments using the cutoff threshold described above.

We compared the proportions of *Caballeronia* positive frass droplets among instars using Fisher’s exact test, and multiple pairwise comparisons among treatments were performed using a pairwise Fisher’s test with a Bonferroni correction (*rstatix* [46]).

## Results

### Experiment 1: Symbiont excretion and bacterial motility

#### Stage-specific symbiont excretion

We found *Caballeronia* present in adult *L. zonatus* feces and largely absent from other life stages (Fisher’s exact test: *P* = 0.0005; Fig. 1). *Caballeronia* was entirely absent from the feces of 2^nd^, 3^rd^, and 4^th^ instar nymphs and was present in very few fecal droplets from 5^th^ instars (∼3%, Fig. 1). In contrast, approximately 80% of adult feces droplets bore *Caballeronia* (Fig. 1).

**Figure 1.**
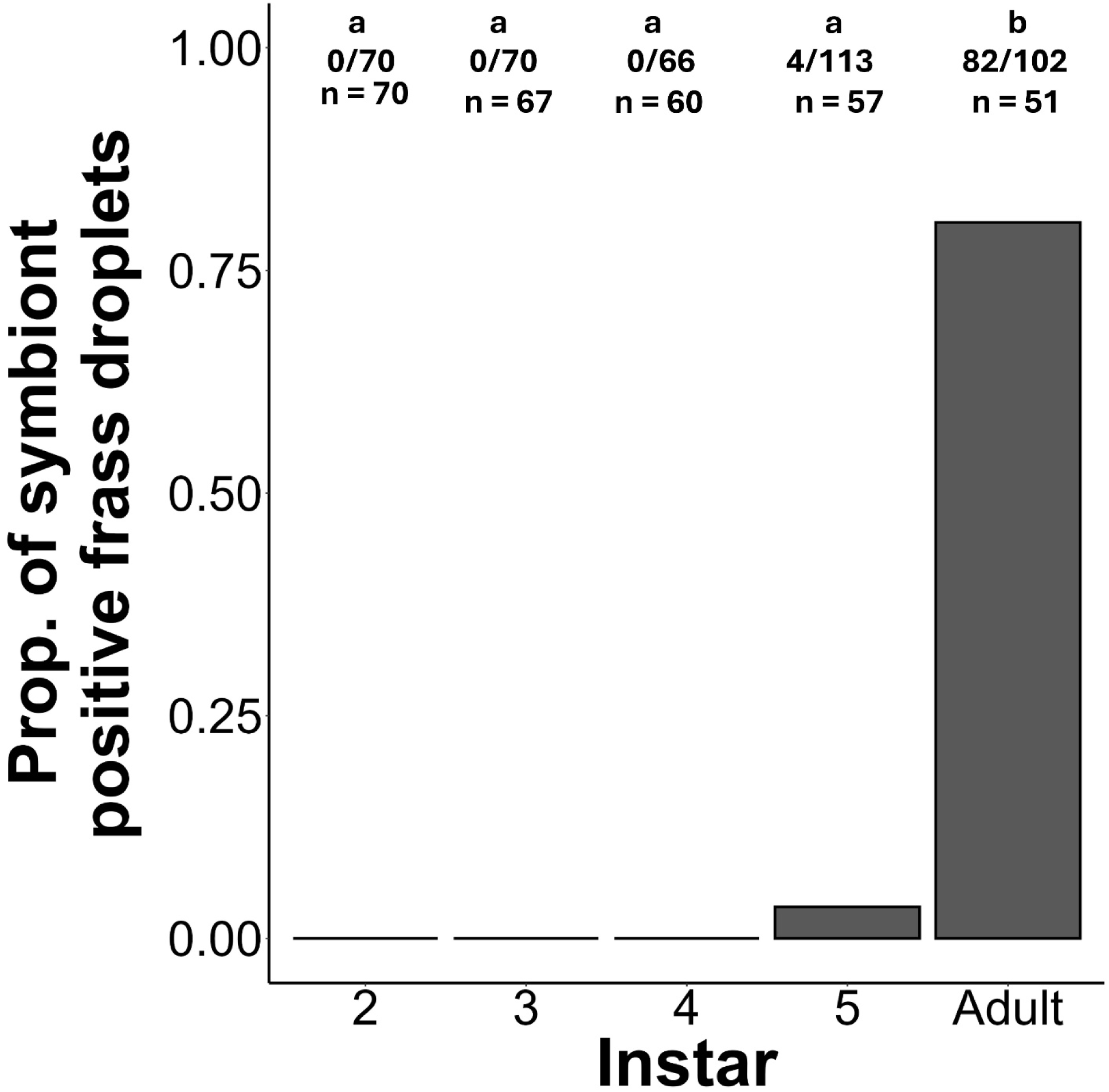
Proportion of fecal droplets containing *Caballeronia* by instar. Letters represent comparisons among treatments using a pairwise Fisher’s exact test with Bonferroni correction for multiple comparisons. Different letters denote treatments that are significantly different (P<0.05). The fractions below represent the number of *Caballeronia* positive frass deposits of the total collected. Sample sizes (the bottom row) are the number of insects that deposited frass that were examined at each life stage.

#### Bacterial motility in culture and in feces

Bacterial motility varied significantly by treatment (Fishers Exact test: *P* = 0.0005). Bacteria were motile in most liquid culture samples examined (85%), but no motile bacteria were observed in either dissected M4 gut sections or in liquid feces (Fig. 2). The proportion of samples with motile bacteria in culture was significantly different from that of samples in the feces treatment (*P* = 2.72×10^−3^) and M4 samples (*P* = 1.94×10^−6^), while motility in feces and from M4 samples were not statistically different from each other (*P* = 1.00; Fig. 2).

**Figure 2.**
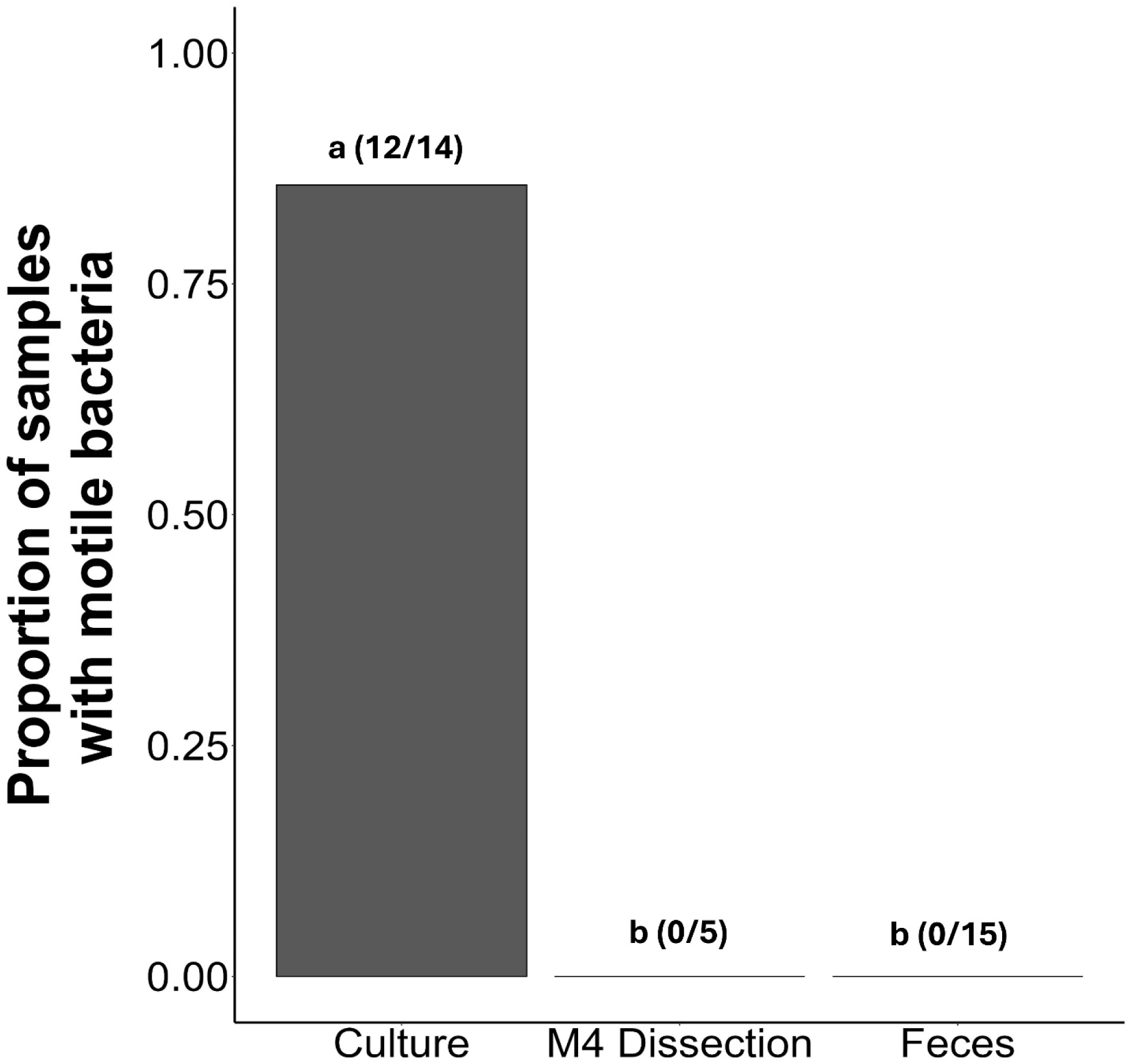
Proportion of video samples with motile bacteria. Five bacteria were assessed in each video sample. Bacteria were considered motile if the average velocity was >4um/s and was independent of fluid flow. Letters represent pairwise comparisons of the three treatment groups. Comparisons were made using a pairwise Fisher’s exact test with Bonferroni correction. The numbers in parentheses are the number of video samples with motile cells out of the total examined.

### Experiment 2: Horizontal transmission of Caballeronia

#### Effect of adult age on transmission

The age of the excreting adult was found to impact only one of the measured variables, adult weight. Adult age of the excreting bug did not significantly alter juvenile acquisition success (LRT = 1.19, *P =* 0.28), development time (LRT = 2.06, *P* = 0.151) or survivorship (LRT = 0.04, *P =* 0.834). For this reason, data from the blocks were pooled and subsequent analyses performed on the full data set. However adult age did have a minor, but significant, impact on adult weight was found to be significantly influenced (*F* = 4.17, *P =* 0.043) and was retained for final analysis.

#### Caballeronia transmission via coprophagy

Symbiont source had no impact on the rate of successful acquisition of *Caballeronia* (LRT = 0.254, *P* = 0.614; Fig 3a). Nymphs were equally likely to acquire the symbiont from feces or from cultured cells. The symbiont-free control bugs were not included in the analysis, since they were not exposed to the symbiont.

**Figure 3.**
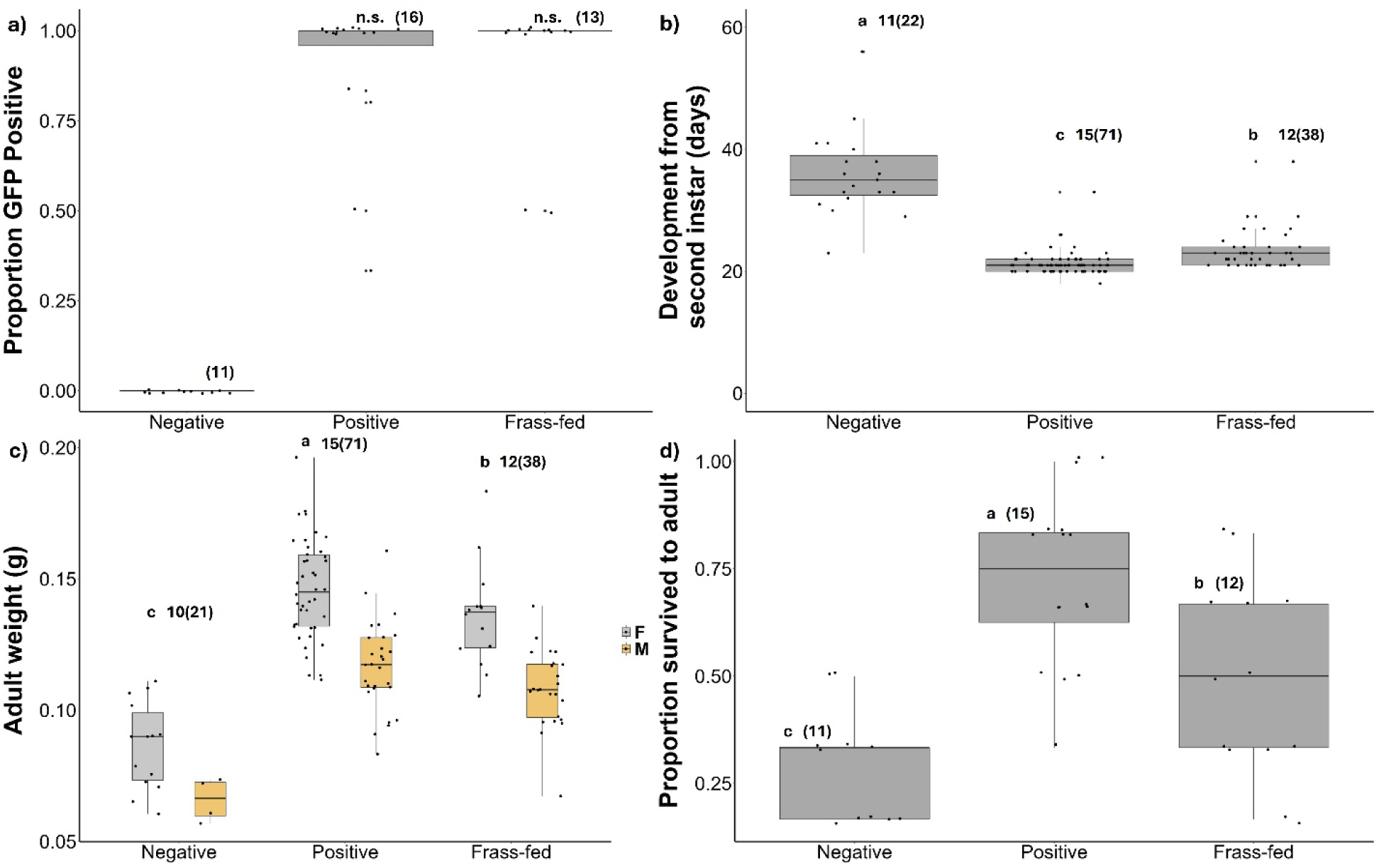
a) Proportion of individuals in each treatment in which the GFP transformed symbiont was detected. b) Development of nymphs in days from second instar to adulthood. c) Individual adult weight by treatment and sex. d) Proportion of individuals that survived to adulthood. Numbers above boxes represent the number of replicates and the numbers within the parentheses are the numbers of individuals measured. Different letters above boxes denote statistically significant differences (P<0.05) representing comparisons among each pair of treatments.

#### Development time

The presence and source of the symbiont had a significant effect on total development time of the insects (LRT = 163.19, *P =* <0.0001; Fig. 3b). Development time from the 2^nd^ instar to adult was shortest in the *Caballeronia* culture-fed treatment (21.2 ± 1.84 days). *Caballeronia* culture-fed individuals developed significantly faster than individuals in the feces-fed treatment (23.6 ± 3.35 days; *z* = 3.65, *P* = 0.0007) and much faster than those in the symbiont-free control (35.8 ± 6.55 days; *z* = -17.82, *P* < 0.0001). The feces-fed treatment bugs also developed significantly faster than the negative control bugs (*z =* -13.49, *P <* 0.0001; Fig. 3b).

#### Adult weight

Adult weight was significantly impacted by sex, with females being approximately 1.2x the size of males (*t* = -8.94, *P =* 5.44×10^−15^). The age group of the defecating adults had a slight but significant impact on the final adult weight of the bugs that ingested the feces at the second instar - older adult defecators led to slightly heavier final adult weights among the bugs that ingested the feces as nymphs (ingestors of feces from younger defecators: 0.121 ± 0.03g, ingestors of feces from mature defecators: 0.123 ± 0.026g; *t* = 4.00, *P* = 0.043). Treatment also had a significant influence on adult weight (LRT = 107.97, *P* = <0.0001; Fig. 3c). Individuals in the *Caballeronia* culture-fed positive control treatment were significantly larger (0.135 ± 0.023 g) than individuals in the feces-fed treatment (0.118 ± 0.021g; *t* = 2.857, *P* = 0.014) and much larger than the aposymbiotic bugs in the symbiont-free control treatment (0.08 ± 0.018g; *t* = 12.73, *P* <0.0001). Insects in the symbiont-free treatment were also significantly smaller than the feces-fed adults (*t* = -9.51, *P <* 0.0001; Fig 3c).

#### Survivorship

Treatment significantly affected survivorship to adult (LRT = 32.25, *P* = 9.93×10^−8^; Fig. 3d). The culture-fed positive control had the greatest proportion of bugs that survived to adulthood (0.75 ± 0.18), significantly more than the feces-fed treatment (0.5 ± 0.23; *z* = 3.19, *P* = 0.004) and the symbiont-free treatment (0.33 ± 0.11; *z* = 5.32, *P* < 0.001). Survival in the symbiont-free treatment was also significantly reduced from the feces-fed treatment (*z* = -2.03, *P =* 0.041; Fig. 3d).

## Discussion

Coprophagy is common among insects, particularly as a route for symbiont transmission [6,47]. While most familiar in social insects such as termites as well as in cockroaches [48–50], some Hemiptera also smear eggs or directly consume conspecific feces [5,6]. However, these examples are often very specialized relationships, with vertical or horizontal transmission serving as the exclusive source of the symbiotic partner(s). Here, we show that coprophagy can serve as an alternate route of symbiont transmission for *L. zonatus. Caballeronia* are present in the feces of *L. zonatus,* with fresh feces containing 2.53×10^5^ CFU/µl (± 3.55×10^5^) of *Caballeronia*. In this way, *L. zonatus* is similar to another coreid bug, *Anasa tristis,* which also shows excretion of *Caballeronia* [28]. Curiously, symbiont excretion in the current study was entirely absent in the 2^nd^ through the 4^th^ instars, and very limited in the 5^th^ instar (4/113 fecal droplets), but present in most adult fecal samples (82/102 droplets). This stage-specific excretion of the symbiont shines light on the potentially interesting gut junction between the M4, brimming with *Caballeronia* cells, and the hindgut from which excretion occurs. In general, *Caballeronia* in the M4 region are refluxed, moving anteriorly, into the M4B where some symbiont cells are digested by the host [51]. In *R. pedestris* the CR re-opens late in the 5^th^ instar, though gut contents from the M3 do not penetrate into the M4 region after this time [51]. It may be that the dynamics of gut flow change near the time of eclosion in *L. zonatus*, allowing movement of *Caballeronia* into the hindgut for eventual release.

We showed that for *L. zonatus* nymphs, as for *A. tristus*, freshly deposited conspecific feces colonized nymphs at equivalent rates to cultured bacteria, indicating these bacteria eventually traverse the CR into the M4. We also observed no significant difference in acquisition success of nymphs presented with feces excreted by adults of different ages, at least up to two weeks after eclosion. Like many members of the coreoid and lygaeoid superfamilies, *L. zonatus* forms large multigenerational aggregations [52–54]. Thus, for these species, feces could serve as a highly concentrated (∼2.5×10^5^ CFU/µl), if potentially ephemeral source of *Caballeronia*. Interestingly, *Caballeronia* is of higher abundance in feces than in cultured *Caballeronia* used for feeding in our experiments, yet we saw decreased survivorship, adult weight, and increased development time in the feces-fed treatment. Of course, cultured *Caballeronia* is not a realistic proxy for cells available in the environment. When acquired from the soil, *Caballeronia* may be flagellate but would be unlikely to be at such high abundance and never in pure culture. It may be that the performance we measured for bugs fed cultured *Caballeronia* was higher than would be typical for bugs acquiring the symbiont from the soil environment in nature. Should that be true, the performance of bugs that acquired the symbiont horizontally from feces might be more comparable to the performance of bugs that acquire it from the natural soil environment. This would be an interesting comparison to make in future studies.

A possible cause of reduced fitness in feces-fed bugs may be the lack of motility of the excreted *Caballeronia.* Within the host*, Caballeronia* is aflagellate and nonmotile, although returns to its flagellate state when transferred to culture media after ∼24hr [14]. We found that *Caballeronia* bacteria are non-motile in fresh *L. zonatus* feces (≤ 24 hours after deposition) as well as in the M4 region, as was shown in *R. pedestris* [14] and in *A. tristis* [40]. As motility is essential for colonization of the bug gut, *Caballeronia* would have to regain motility in order to transit the constricted region. The non-motile *Caballeronia* derived from feces must undergo the necessary transition to the flagellate state to cross into the M4, and this delay between ingestion and colonization of the M4 may result in fitness costs [35]. Other physiological changes or stresses associated with transit through the bug hindgut or with other chemical components of feces may also delay colonization. It could also be that while cell number is initially higher in feces than in culture, the environmental conditions rapidly erode the number of viable *Caballeronia* cells, thus reducing the founding number of colonizing cells in the M4, and potentially delaying proliferation within the M4 and the delivery of nutrients to the growing bug. Previously we showed that when confined to the canopy, the arboreal *L. zonatus* rarely acquires *Caballeronia,* even when nymphs were confined with an adult bug [34]. We might hypothesize that *Caballeronia* survivorship is relatively low on orchard canopy plant surfaces which are potentially subject to high temperatures, low humidity, and harmful UV radiation. In addition, *L. zonatus* population size may be a factor in the likelihood of horizontal transmission, as it may require high densities of excreting adult bugs for live *Caballeronia* to be accessible to nymphs.

The mechanism underlying the fitness differences associated with the source of the symbiont requires further investigation, but whatever the mechanism, the costs could multiply in a more complex environment where *Caballeronia* cells of different strains compete to colonize the M4. *Caballeronia* strain dominance within the gut is driven by priority effects in colonization as well as competition within the M4 [19,55]. When multiple *Caballeronia* strains or species have been ingested, the sooner a strain enters the M4, the higher the likelihood it will proliferate there. Aflagellate *Caballeronia* are likely to be at a disadvantage against an already flagellate strain or species ingested at the same time.

Environmental acquisition is likely to explain much of the heterogeneity in *Caballeronia* strains seen in these insects. Soil is inhabited by diverse *Caballeronia* communities [19,56,57] and those communities are likely structured by biotic factors as well as heterogeneity in aridity, temperature, pH, pesticide use and nutrient availability [58–60]. When a bug acquires its *Caballeronia* from the soil, numerous strains may compete and colonize the M4, although competitive interactions are likely to reduce the community complexity in the gut relative to the soil [19]. Multiple infections in individuals are common in *L. zonatus* as well as *R. pedestris* [17,23]. Should feces be the primary symbiont source, however, we would anticipate that symbiont diversity would decline over generations as some strains dominate competitive interactions within the bug, with single infections becoming more common. We do not see this pattern in most bugs that host *Caballeronia* however, as multiple infections appear frequently in nature [19,34,36,56,61,62].

In this context, horizontal transmission via coprophagy is likely to play an interesting if potentially limited role in the maintenance of this symbiosis (57–59). Acquisition of *Caballeronia* symbionts from complex and dynamic soil communities is likely to maintain the large and multifunctional genomes characteristic of free-living bacteria, while horizontal transmission may play a less frequent but important role for the offspring of adult bugs colonizing areas where the soil is depauperate in *Caballeronia.* Further, excreted *Caballeronia* strains are, by definition, successful in providing sufficient nutrition for growth of the adult bug that excreted them, and in this way adults may enrich local environments with strains that are beneficial, locally adapted and competitive in the bug gut. In general, one might expect selection on hosts to increase the likelihood of acquisition of a critical partner via vertical transmission, though the long association of several families of coreoid and lygaeoid bugs with environmentally acquired *Caballeronia* appears an exception to this idea [20]. For these bugs instead, utilizing an ‘unfaithful’ [63] or a hybrid transmission strategy, one that incorporates both horizontal transmission and environmental acquisition, may provide resilience to change and adaptive solutions for novel environments.

## Acknowledgements

We would like to thank Jackson Schwenn for assistance with dissections and imaging of gut tissue.

## Author Contributions

Liam T. Sullivan: Writing – original draft, Methodology, Data curation, Formal Analysis; Suzanne E. Kelly: Conceptualization, Methodology; Martha S. Hunter: Conceptualization, Supervision, Writing – review & editing

